# Multi-trait random regression models increase genomic prediction accuracy for a temporal physiological trait derived from high-throughput phenotyping

**DOI:** 10.1101/772038

**Authors:** Toshimi Baba, Mehdi Momen, Malachy T. Campbell, Harkamal Walia, Gota Morota

## Abstract

Random regression models (RRM) are used extensively for genomic inference and prediction of time-valued traits in animal breeding, but only recently have been used in plant systems. High-throughput phenotyping (HTP) platforms provide a powerful means to collect high-dimensional phenotypes throughout the growing season for large populations. However, to date, selection of an appropriate statistical genomic framework to integrate multiple temporal traits for genomic prediction in plants remains unexplored. Here, we demonstrate the utility of a multi-trait RRM (MT-RRM) for genomic prediction of daily water usage (WU) in rice (*Oryza sativa*) through joint modeling with shoot biomass (projected shoot area, PSA). Three hundred and fifty-seven accessions were phenotyped daily for WU and PSA over 20 days using a greenhouse-based HTP platform. MT-RRMs that modeled additive genetic and permanent environmental effects for both traits using quadratic Legendre polynomials were used to assess genomic correlations between traits and genomic prediction for WU. Predictive abilities of the MT-RRMs were assessed using two cross-validation (CV) scenarios. The first scenario was designed to predict genetic values for WU at all time points for a set of accessions with unobserved WU. The second scenario was designed to forecast future genetic values for WU for a panel of known accessions with records for WU at earlier time periods. In each scenario we evaluated two MT-RRMs in which PSA records were absent or available for time points in the testing population. Moderate to strong genomic correlations between WU and PSA were observed across the days of imaging (0.29-0.87). In both CV scenarios, MT-RRMs showed better predictive abilities compared to single-trait RRM, and prediction accuracies were greatly improved when PSA records were available for the testing population. In summary, these frameworks provide an effective approach to predict temporal physiological traits that are difficult or expensive to quantify in large populations.

## Background

High-throughput phenotyping (HTP) is an innovative tool in plant breeding. HTP provides precise and non-destructive estimation of multiple complex traits that describe growth and development (e.g., height, biomass, and flowering time) or environmental responses (e.g., chlorophyll fluorescence, canopy temperature, and water content) using non-destructive image-based phenotyping (Araus et al., 2018; Morota et al., 2019). These HTP data mitigate extensive costs associated with manual phenotyping, and can be used to better capture the plant’s phenome. In the context of plant breeding and genetics, these data can be used to improve the prediction of breeding values for a target trait of interest, thereby improving the accuracy of selection, as well as provide insights into how secondary traits influence a trait of interest (Araus et al., 2018; Morota et al., 2019; Voss-Fels et al., 2019).

For many genome-enabled breeding programs, developing phenotyping and statistical approaches to improve prediction of breeding values and accelerate selection is the primary objective (Campbell et al., 2018; Juliana et al., 2019; Voss-Fels et al., 2019). In many breeding programs, the agronomic value of breeding materials is evaluated using multiple traits. These traits are often correlated at the genetic level. One standard approach for predicting breeding values is to jointly fit all phenotypes in a single model using a multi-trait (MT) approaches (Kadarmideen et al., 2003). These approaches capture the genetic covariances between traits, and have been shown to improve the prediction of breeding values compared to single trait approaches for phenotypes with limited records or low heritability (Calus and Veerkamp, 2011; Jia and Jannink, 2012; Guo et al., 2014). Thus, the MT framework can be particularly advantageous when the target trait has low heritability, but is correlated with a more heritable trait; or when the trait of interest is difficult or costly to evaluate and only incomplete data can be collected, and the trait of interest is correlated with a trait that is easier and cheaper to evaluate. Thus, in the context of HTP, MT genomic prediction approaches can accommodate the high-dimensional multi-trait data generated by these platforms. Moreover, secondary phenotypes recorded with HTP can be included in the prediction framework to improve prediction of a target trait such as yield. These applications have been shown in a recent study by (Sun et al., 2017).

While several studies have highlighted the advantages of MT frameworks for genomic prediction, HTP-derived MT data often introduce an additional level of complexity-the time axis. The standard MT framework may not be appropriate in cases where multiple phenotypes are recorded at regular intervals throughout the growing season or for the duration of the experiment. While MT frameworks can be fit to these data, the assumptions of the MT framework bring to question whether the conventional MT model should be used. For instance, one assumption is that each phenotype in the MT model is finite characteristic (Kirkpatrick et al., 1990). While this is certainly true for two phenotypes such as yield or protein content, this is certainly not the case for a phenotype recorded at two time points (Kirkpatrick et al., 1990). Temporal phenotypes are infinite-dimensional traits, meaning that although there are only records for discrete time intervals, we expect that the phenotype will vary continuously with time between the two intervals. With these data, a more appropriate solution is to treat the temporal phenotypes as continuous characteristics and perform genetic analyses using random regression models (RRM).

RRM model the covariance between time points as a continuous function of time (Mrode, 2014). While several covariance functions can be utilized, Legendre polynomials or B-splines are routinely used. The use of orthogonal Legendre polynomials in RRM offers numerical stability by reducing correlation between random regression coefficients and computing error (Schaeffer, 2004). With RRM, temporal phenotypes are partitioned into genetic, permanent environmental effects, and residuals (Mrode, 2014). With repeated measurements, it is assumed that there is additional resemblance between records of an individual due to environmental factors or circumstances that affect the records of the individual permanently (Mrode, 2014). Thus, the random permanent environment term captures this non-genetic resemblance between time points. Covariance functions are used to model both genetic and permanent environmental effects (Kirkpatrick et al., 1990; Schaeffer and Dekkers, 1994; Meyer and Hill, 1997; Schaeffer, 2004). Thus, the RRM prediction framework provides solutions for random regression coefficients for random effects. Given coefficients for random genetic effects, the genetic values at any time point can be easily calculated. Recently, RRM have been used for genomic analyses of longitudinal image-based HTP traits in plants (Campbell et al., 2018, 2019; Momen et al., 2019a). The ability of these frameworks to forecast future phenotypes using the records at earlier time has been shown by Campbell et al. (2018) and Momen et al. (2019a) based on a digital metric for shoot biomass, known as projected shoot area (PSA). PSA is a digital metric derived from images taken of each plant and is highly correlated with destructive measures of shoot biomass (Golzarian et al., 2011; Berger et al., 2010; Campbell et al., 2015).

However, given the capability of HTP to collect multiple temporal phenotypes, one unresolved question in plant breeding is how to jointly model multiple temporal phenotypes. To address this, we aimed to integrate the RRM framework for temporal traits into a MT model. We utilized a data set in which PSA and water use (WU) was recorded daily over a period of 20 days. The aim of the study was to evaluate the ability of multi-trait random regression model (MT-RRM) and a single-trait random regression model (ST-RRM) to predict WU by borrowing information from PSA. The rationale is that WU is much more difficult to evaluate in most studies compared to PSA and is likely to be more influenced by environmental effects, and thus have lower heritability compared to shoot biomass. The models were compared using several cross-validation (CV) scenarios.

## Materials and Methods

### Plant materials and greenhouse conditions

This study utilized HTP records from 378 of the 432 accessions of rice (*Oryza sativa*) diversity panel 1 (RDP1) (Zhao et al., 2011). Sixty four accessions were excluded due to lack of seed availability or poor germination. Seeds were treated with Thiram fungicide and germinated on moist paper towels in plastic boxes for three days. Three uniformly germinated seedlings were selected for each accession and transplanted to pots (150mm diameter x 200 mm height) filled with 2.5 kg of UC Mix. The plants were grown in saturated soil on greenhouse benches prior to phenotyping.

Plants were thinned to one seedling per pot seven days after transplant (DAT), and two layers of blue mesh were placed on top of the pots to reduce evaporation. The plants were loaded on to the imaging system at 13 DAT. The automated phenotyping system was set to maintain all plants at 90% field capacity. The experiment followed a partially replicated design (Cullis et al., 2006). The p-rep design was modified to accommodate the two water treatments (control and drought conditions) and allow comparison of treatments within each accession. Each accession was assigned to two consecutive pots, and the water treatments were randomly assigned to each pot. Each experiment consisted of 378 accessions from RDP1 and was repeated three times from February to April 2016. The accessions were distributed across 432 pots positioned across 24 lanes (18 plants/pots in each lane). These 432 pots belonged to 378 accessions, of which 54 had more than one replicate in each experiment. The same 54 accessions were replicated twice in each experiment. Of these 378 accessions, 357 accessions had genotypic data. All experiments were conducted at the Plant Accelerator Australian Plant Phenomics Facility, at the University of Adelaide, SA, Australia.

### Phenotypic data

Beginning at 13 DAT all plants were phenotyped daily for shoot biomass and WU using the automated greenhouse system, and each plant was imaged daily over a period of 20 days using a visible (red-green-blue / RGB) camera (Basler Pilot piA240012 gc, Ahrensburg, Germany). For each plant, three images were taken in each recording day: two side-view angles separated by 90 degree and a single top view. Plant pixels were extracted from RGB images using the LemnaGrid software, and the plant pixels from the three images were summed to obtain a digital measure of shoot biomass. We refer to this metric as PSA. Several studies have shown this to be an accurate proxy for shoot biomass (Golzarian et al., 2011; Campbell et al., 2015; Knecht et al., 2016).

After imaging, each plant was watered to a predefined weight to maintain 90% field capacity. The automated watering system collects the start weight, final weight and amount of water that was added for each pot. Thus, from these data we can estimate the amount of water that lost by evapotranspiration each day. WU was calculated as *WU*_*t*_ = *Potwt*_*t*−1_ − *Potwt*_*t*_. Where *Potwt*_*t*−1_ is the weight of the pot after watering on the previous day, and *Potwt*_*t*_ is the weight of the pot on the current day prior to watering (Momen et al., 2019b).

In this study, we used observations collected in the control condition. Best linear unbiased estimators (BLUE) were obtained for each accession and day using the following model

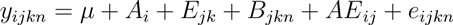

where *µ* is the overall mean, *A_i_* is the effect of the *i^th^* accession, *E_jk_* is the effect of the *j^th^* experiment in the *k^th^* replicate, *B_jkn_* is the block effect of the *n^th^* smart house in the *j^th^* experiment and the *k^th^* replicate, *AE_ij_* is the interaction of accession and experiment. All the effects, except *A_i_* were considered random.

### Genotypic data

All accessions were genotyped with a 44,000 single nucleotide polymorphisms (SNPs) array (Zhao et al., 2011). Genotypic data regarding the rice accessions can be downloaded from the rice diversity panel website (http://www.ricediversity.org/). SNPs with call rate ≤ 0.95 and minor allele frequency ≤ 0.05 were removed. Missing genotypes were imputed using Beagle software version 3.3.2 (Browning and Browning, 2007) following Momen et al. (2019b). A total of 34, 993 SNPs remained for downstream analyses.

### Single-trait random regression model

Campbell et al. (2018) and Momen et al. (2019a) have applied ST-RRM for PSA. In this study, a similar statistical model was used to model WU. The model is given by

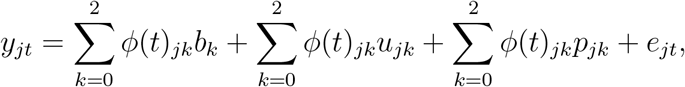

where *y*_*jt*_ is the BLUE of *j*th accession for WU at time point *t*, *b*_*k*_ is the *k*th fixed Legendre regression coefficients for overall mean, *u*_*jk*_ is the *k*th random regression coefficients for additive genetic effect, *p*_*jk*_ is the *k*th random regression coefficients for permanent environmental effect, *e*_*jt*_ is the vector of residuals, and *φ*(*t*)_*jk*_ is a time covariate coefficient defined by a *k*th Legendre polynomial evaluated at time point *t* belonging to the *j*th accession. The permanent environmental effect captures constant environmental factors that affect the successive records of an accession throughout the time course (Mrode, 2014). We set quadratic Legendre polynomials of all the effects, based on the results of Momen et al. (2019a) which investigated the prediction accuracy of PSA using ST-RRM. The first order of the Legendre polynomial (i.e., an intercept) was standardized to 1 (Gengler et al., 1999).

In matrix notation, the model is given by

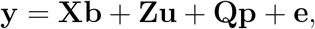

where **y** is the vector of observations for WU, **b** is the vector of fixed effect, **u** is the vector of random additive genetic effect, **p** is the vector of random permanent environmental effect, **e** is the vector of random residual effect, and **X**, **Z**, and **Q** are corresponding incidence matrices. The covariance structures were defined as the following.

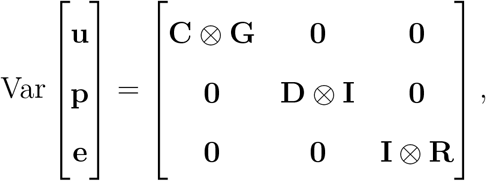

where **G** is a genomic relationship matrix calculated by **WW**′/*m* according to VanRaden (2008), **W** is a centered and scaled matrix, *m* is the number of SNPs, **I** is an identity matrix, **C** and **D** are covariance matrices of additive genetic and permanent environmental effects, **R** is a diagonal matrix of heterogeneous residual variance at each time period, and ⊗ is the Kronecker product. The covariance matrices **C** and **D** are defined as follows.

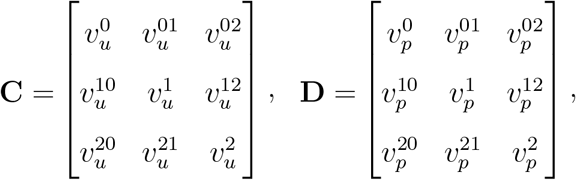

where 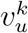 and 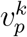 are the variance components of *k*th order random regression coefficients for additive genetic and permanent environment effects, respectively, and 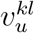 and 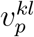 are the covariances between *k*th and *l*th order random regression coefficients for additive genetic and permanent environmental effects, respectively.

### Multi-trait random regression model

For MT-RRM, the ST-RRM for WU described above is expanded to include PSA information as follows.

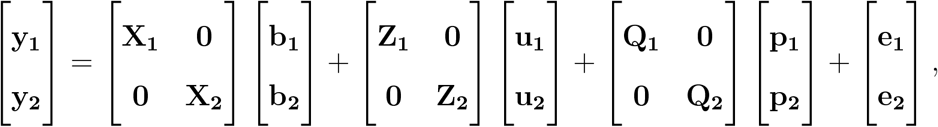

where subscripts 1 and 2 refer to WU and PSA, respectively. The covariance structures of **C** and **D** were also expanded as follows.

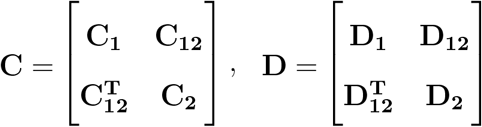

where **C**_**1**_ and **C**_**2**_ (**D**_**1**_ and **D**_**2**_) are 3×3 variance-covariance submatrices of random regression coefficients for each trait and **C**_**12**_ (**D**_**12**_) is a 3 × 3 covariance submatrix of random regression coefficients between the traits. Thus, the whole **C** and **D** matrices take the form

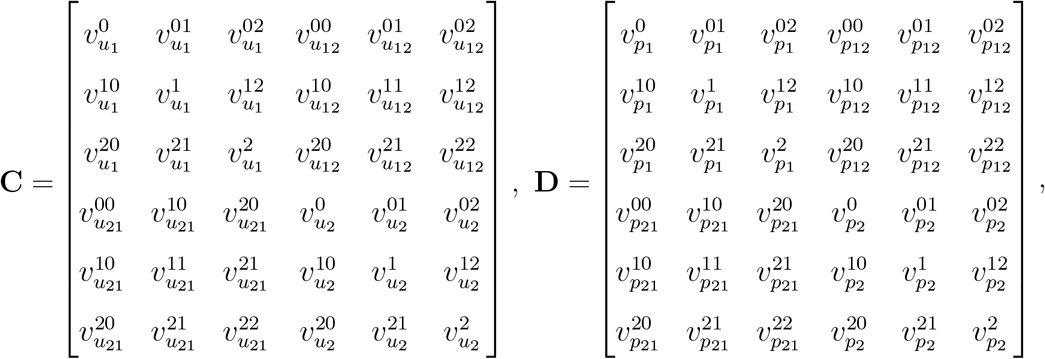

where 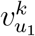 and 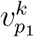 (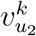 and 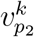) are variance components of *k*th order random regression coefficients for additive genetic and permanent environment terms for WU (PSA), 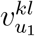 and 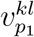 (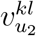 and 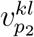) are covariances between *k*th and *l*th order random regression coefficients for additive genetic or permanent environmental effects within WU (PSA), and 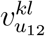 and 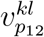 are covariances between *k*th and *l*th order random regression coefficients for additive genetic and permanent environmental effects between WU and PSA, respectively. As with ST-RRM, we assumed the residual variance for each day of imaging was unique. Thus, a heterogeneous residual variance structure was used for MT-RRM. The matrix of residual variance at time 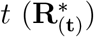 is presented as:

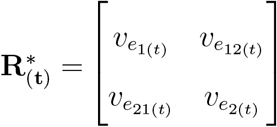

where 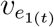 and 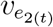 are residual variances for WU and PSA, respectively, and 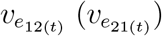 is the residual covariance between WU and PSA at time point *t*.

### Estimation of genomic correlation at each time point

Genomic correlation between WU and PSA at each time point from MT-RRM was computed as follows.

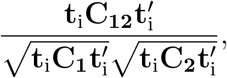

where **t**_**i**_ = *φ*_*ik*_ is the *i*th row vector of the 20 × 3 basis function matrix (**Φ**) with the *k*th order of fit (Mrode, 2014). Here, **Φ** is given as **MΛ**, where **M** is a matrix of second order polynomials of standardized time values and **Λ** is a matrix of coefficients for a second order Legendre polynomial (Kirkpatrick et al., 1990). We used the GIBBS3F90 program to estimate genetic parameters (Misztal et al., 2002). The GIBBS3F90 program solves mixed model equations in the Bayesian framework by assuming heterogeneous residual variances.

### Cross-validation scenarios

We investigated the prediction performance of genetic values for WU from RRM using two CV scenarios as shown in Figure 1. For each CV scenario, we compared three models as described below.

**Figure 1:**
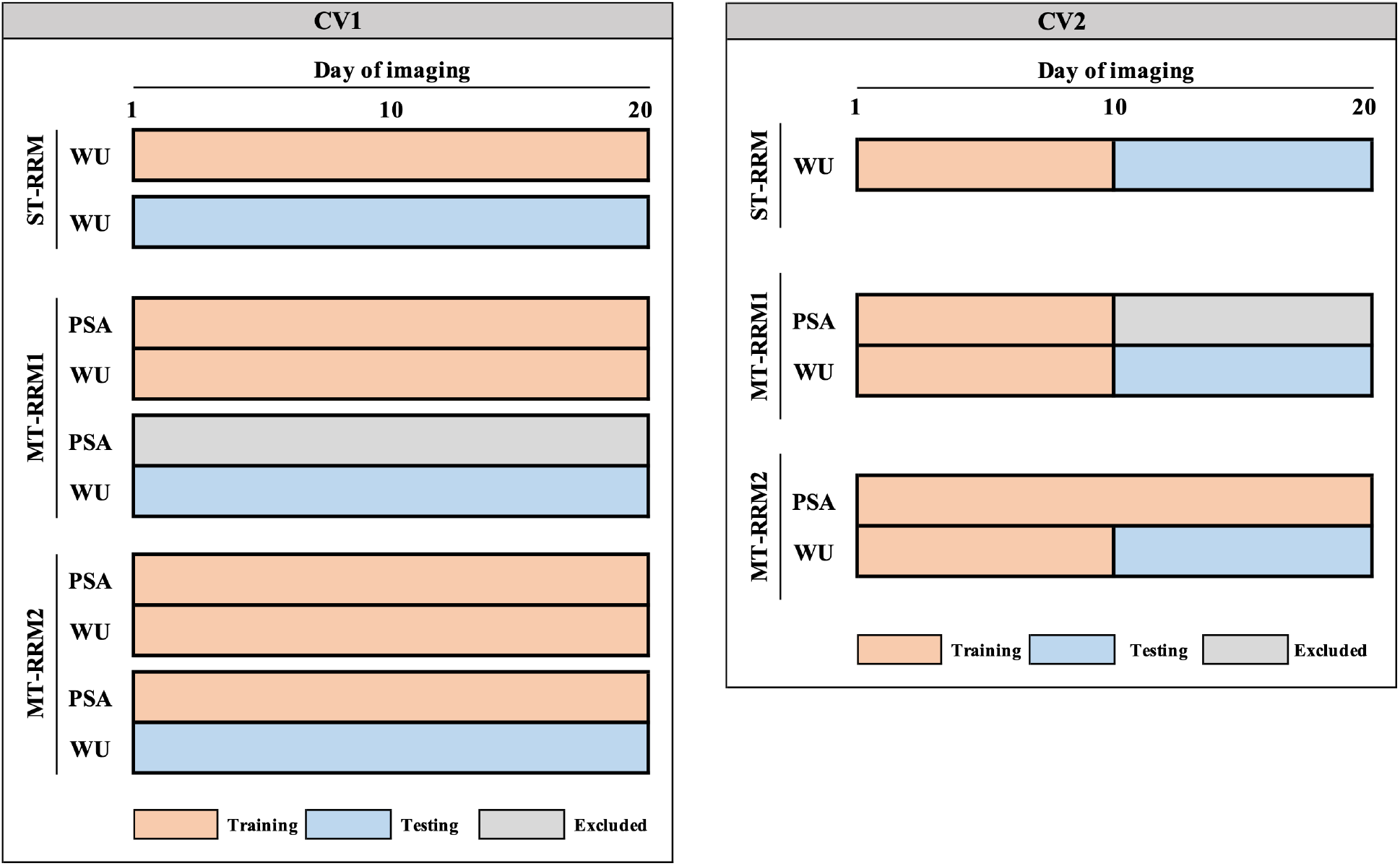
Two scenarios of cross-validation (CV) designed to investigate prediction accuracy of water use (WU) from single- and multi-trait random regression models (ST-RRM, MT-RRM1, and MT-RRM2). CV1: Prediction of WU for a set of 112 accessions without records on WU using 245 training accessions. ST-RRM: single-trait random regression model using WU of training accessions; MT-RRM1: multi-trait random regression model using WU and projected shoot area (PSA) of training accessions; MT-RRM2: multi-trait random regression model using WU and PSA of training accessions as well as PSA of testing accessions. CV2: Forecast future genetic values of WU belonging to 245 known accessions from records at earlier time periods. ST-RRM: single-trait random regression model for WU using the first 10 time points in training accessions; MT-RRM1: multi-trait random regression model for WU using the first 10 time points of WU and PSA information in training accessions; MT-RRM2: multi-trait random regression model for WU using WU from 1 to 10 time periods and PSA at all the time periods in training accessions.

#### CV1

The objective of this scenario was to assess the ability of ST-RRM and MT-RRM to predict WU for a set of new accessions without records on WU. To this end, the accessions were split into testing and training sets with 245 accessions allocated to the training set and 112 allocated to the testing set. First, we fitted ST-RRM using genomic and phenotypic data on the training subset and the genetic values of WU were predicted for all accessions in the testing set. This ST-RRM served as a baseline to evaluate MT-RRM. We evaluated two different types for MT-RRM. The first MT-RRM (MT-RRM1) can be thought of as a conventional genomic prediction application in which a model is fitted using a training population that has genomic data and phenotypic records for both traits. This model is used to predict genomic values for WU in a testing population that has genotypic data, but no records for either trait. In the second MT-RRM (MT-RRM2), complete PSA and WU records were available for the training population, while only PSA phenotypes were available for the testing population. The rationale for this scenario is that it is often much easier to obtain non-destructive measurements for shoot biomass compared to WU. Thus, this can be thought of as a case in which a portion of the population has incomplete data.

Genetic values of testing individuals for WU at time *t* from ST-RRM and MT-RRM1 were calculated by 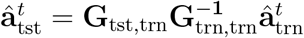, where **G**_tst,trn_ is the genomic relationship matrix between testing and training individuals, 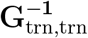 is the inverse of genomic relationship matrix of training individuals, and 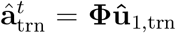 is the vector of genetic values at time *t* (Momen et al., 2019a). On the other hand, the genetic values of testing individuals for WU from MT-RRM2 can be directly obtained from best linear unbiased prediction (BLUP) solutions because the model included the genomic relationship matrix of all accessions by fitting PSA phenotypes for the testing individuals. Thus, the genetic values of WU for the testing individuals at time *t* were computed by 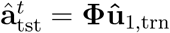.

#### CV2

This cross-validation was designed to evaluate the ability of the MT-RRM and STRRM to predict genetic values of WU at future time points. Thus, it can be thought of as a forecasting approach. The training dataset consisted of phenotypic records for 245 randomly selected for the first 10 days of imaging. The models were used to predict genetic values for days 11 to 20. As with the first scenario, we assessed their genomic predictions by using ST-RRM and two kinds of MT-RRM from the training data. MT-RRM1 used records of WU and PSA from day 1 to day 10 of imaging as training data, while MT-RRM2 used WU records from the first 10 days of imaging and PSA values from 1 to 20 to train the model. We computed the genetic values of WU at day 11 to 20 as **Φ**_11:20_**û**_1,trn_ where **Φ**_11:20_ is the basis function matrix at 11 to 20 days and **û**_1,trn_ is the vector of random additive genetic effect for WU of testing individuals.

To assess prediction accuracy, Pearson correlation was calculated between predicted genetic values and BLUE of WU at each time point in the testing population. Each CV scenario was repeated 10 times. We used the GIBBS3F90 program with a fixed variance option to perform genomic prediction in all the CV scenarios. We estimated variance components in the training set and genetic values were predicted in the testing set condition on the estimated variance components.

## Results

### Assessing temporal water use and shoot biomass trajectories in rice

To assess the temporal relationships between shoot biomass production and WU, a panel of 357 rice accessions was phenotyped over a period of 20 days using a non-destructive image-based phenotyping platform. This system provides a means to non-destructively assess plant growth and morphology, and allows WU to be assessed throughout the duration of the experiment (Berger et al., 2010; Campbell et al., 2015; Fahlgren et al., 2015; Feldman et al., 2018).

Figure 2 shows a boxplot of the BLUE after an adjustment by fixed effects for WU over 20 time days of imaging. WU exhibited an exponential trend over the 20 days of imaging and closely followed the temporal patterns exhibited by PSA.

**Figure 2:**
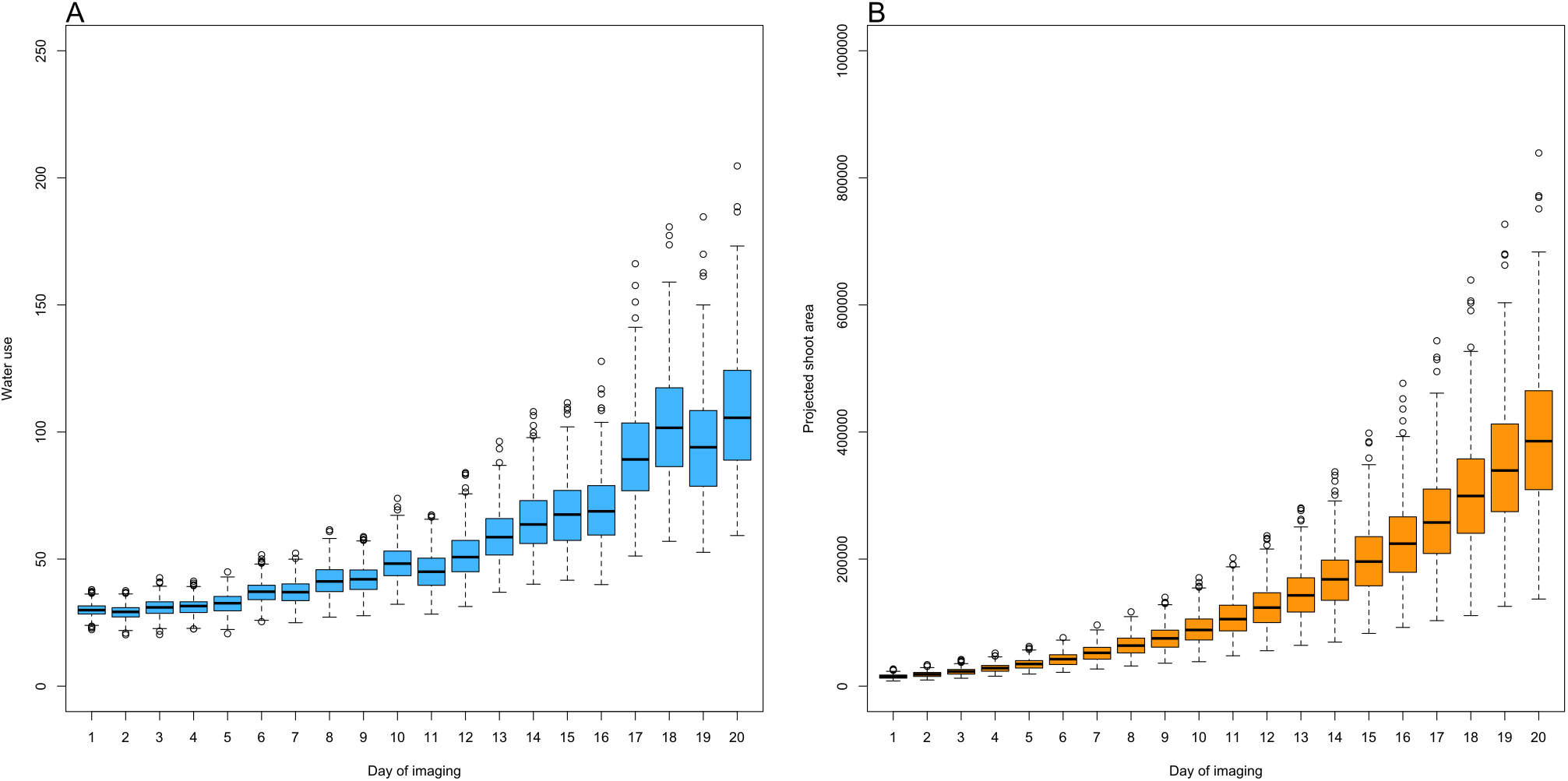
A boxplot of best linear unbiased estimator for water use (A) and projected shoot area (B) over 20 days of imaging.

### Joint analysis of WU and PSA reveals shared additive genetic effects between traits

Genetic architectures of WU and PSA were dissected by estimating the proportion of captured additive genetic variances across 20 days of imaging using a ST-RRM. The RRM included a fixed second order Legendre polynomial to capture the overall mean trajectories for each trait, and additive genetic and permanent environmental effects were modeled using a second order Legendre polynomial. Figure 3 shows that PSA exhibited considerably higher narrow-sense heritability (*h*^2^) compared to WU. For instance, *h*^2^ ranged from 0.48 to 0.82 for PSA, while the values ranged from 0.20 to 0.73 for WU. We observed an increasing trend over time with the lowest value observed on day 1 of imaging and the highest value observed on day 19. PSA on the other hand showed the lowest *h*^2^ values on day 1, but it quickly increased and reached somewhat of a plateau from day 3 to 16. After day 16, *h*^2^ slowly declined. Collectively, these results indicate that both traits are influenced by additive genetic effects, and these effects vary throughout time. However, phenotypic variance is less influenced by non-genetic effects for PSA compared to WU. Moreover, additive genetic effects for WU show greater temporal variability compared to PSA.

**Figure 3:**
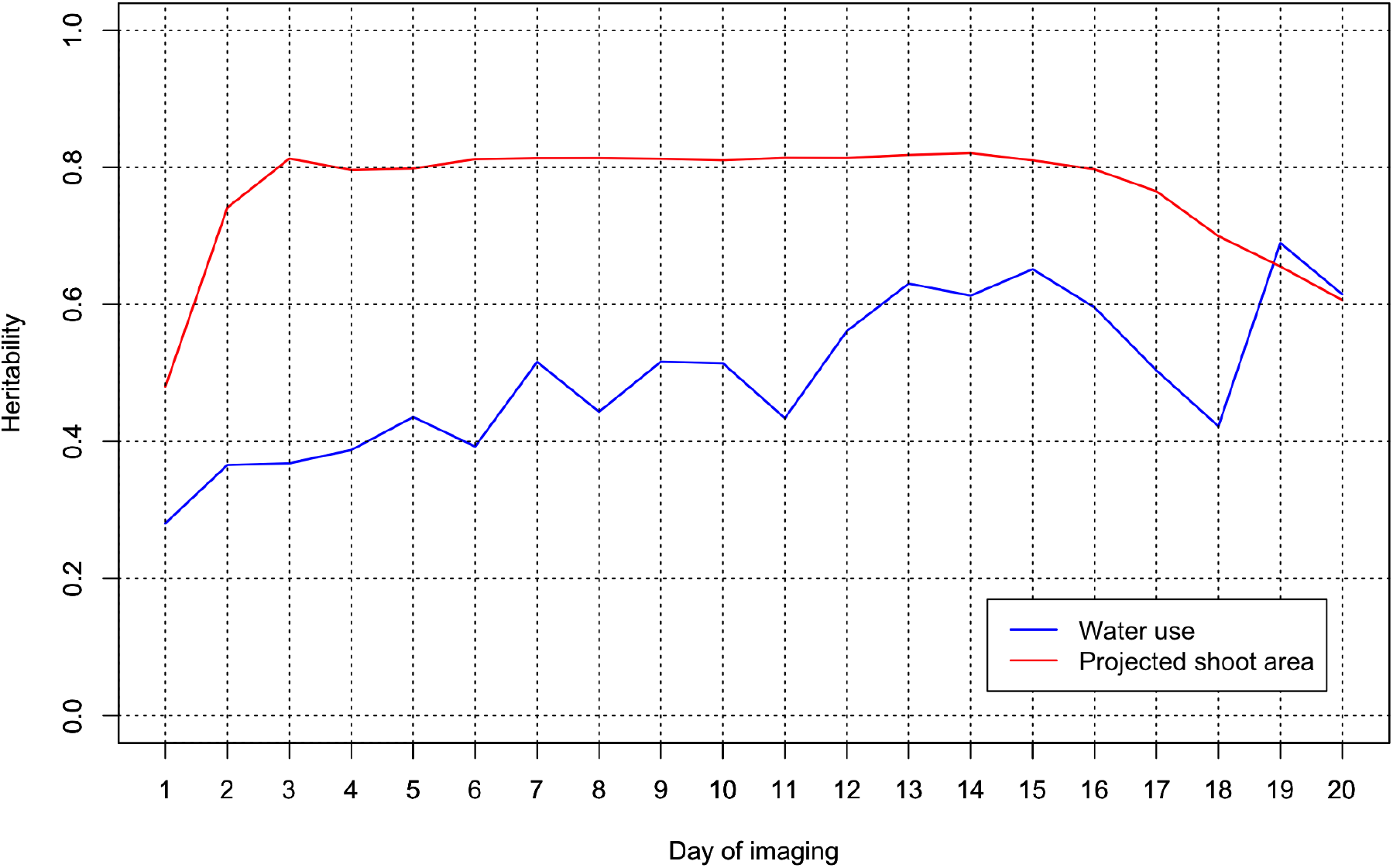
Heritability for water use and projected shoot area over 20 days of imaging using a single-trait random regression model.

To investigate genetic relationship between WU and PSA, we estimated genomic correlation at each time point. The MT-RRM used a second order polynomial to model the overall mean trends for each trait, as well as the additive genetic and permanent environmental effects for both traits. The genomic correlation between WU and PSA from MT-RRM is shown in Figure 4. A moderate to strong positive genomic correlation between WU and PSA was observed over 20 time periods. On average, the genomic correlation across all time periods was 0.78. The genomic correlation was low for the first time point, but quickly increased until the fifth day of imaging. From day 5 to the final day of imaging, genomic correlation showed a slight increasing trend. Genomic correlation ranged from 0.29 to 0.87, with the highest value observed on the last 12 days of imaging. These results indicate that WU and PSA share similarity at the genetic level.

**Figure 4:**
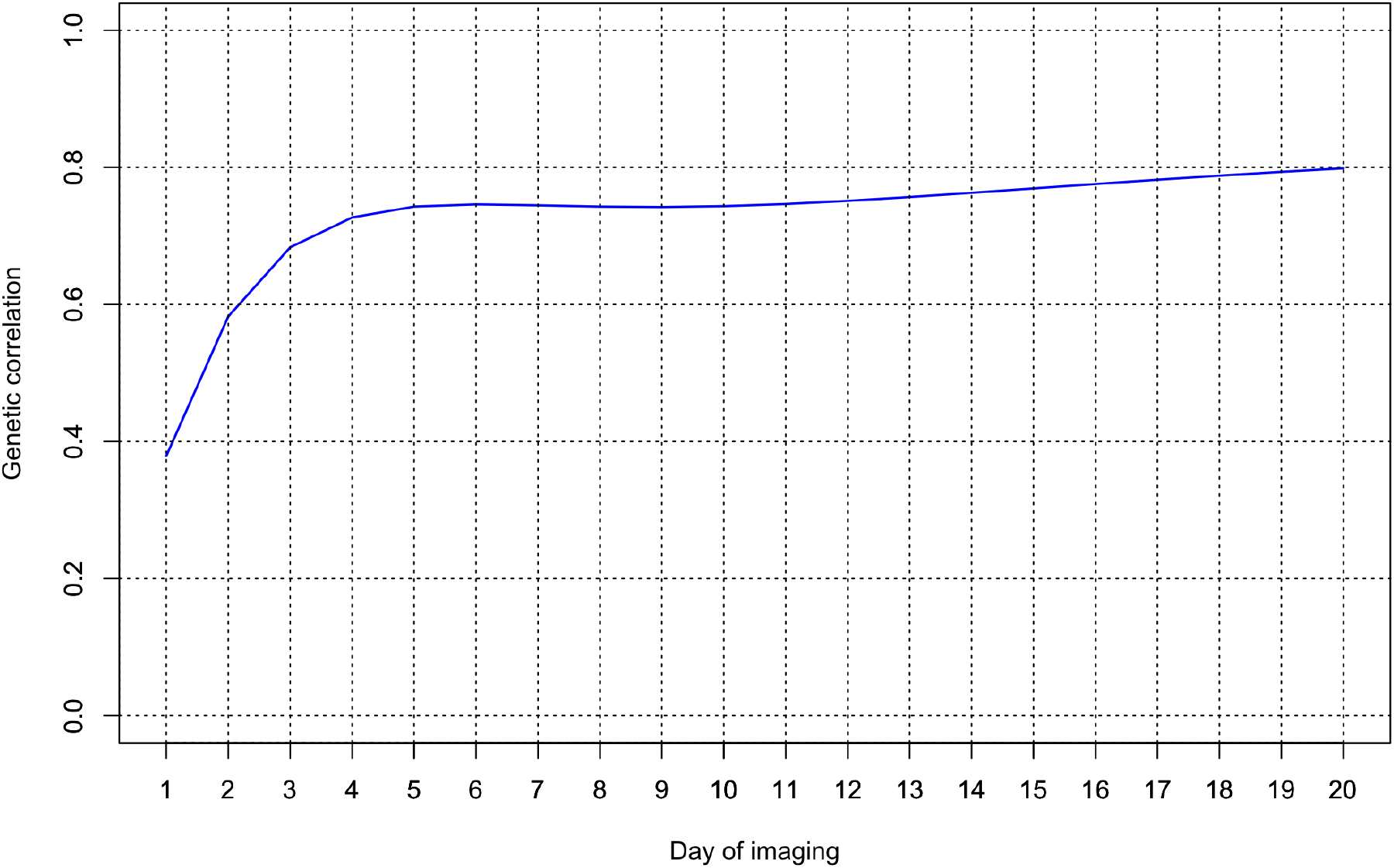
Genomic correlation between water use and projected shoot area over 20 days of imaging using a multi-trait random regression model.

### Predictive assessment using RRM by two CV scenarios

We next sought to evaluate the predictive performance of MT-RRM to predict genetic values for WU. To this end, we employed two CV scenarios. The first is similar to a conventional genomic prediction application in which the objective is to predict genetic values for a set of individuals without phenotypic records. We used two different testing populations. The first consists a set of 112 randomly selected accessions that have no phenotypic records for PSA and WU. The second consists of a set of 112 accessions that have phenotypic records for PSA, but lack records for WU. Thus, the latter scenario can be thought of as a case where a subset of the population has incomplete data. The predictive ability of MT-RRM1 and MT-RRM2 was compared to a ST-RRM in which the model fitted using WU values for 245 accessions and is used to predict genetic values for the remaining 112 accessions. In all cases, the predictive ability was measured as the correlation between predicted genetic values and BLUE in the testing set at each time point.

The prediction accuracy for CV1 is shown in Figure 5. An increasing trend in prediction accuracy over time for all models was observed in CV1. Prediction accuracy increased quickly from day 1 to 13, and eventually plateaued from days 14 to 20. MT-RRM2 showed the highest prediction accuracy of all models. On average the predictive ability for MT-RRM2 was 0.74, while for MT-RRM1 and ST-RRM the prediction accuracies were 0.53 and 0.46, respectively. These results indicate that the predictive ability can be improved using a MT-RRM when records for one trait are available for individuals in the testing population. The predictive ability for MT-RRM1 was similar to ST-RRM during the first five time points, but slightly increased with 0.06 to 0.09 relative to ST-RRM from day 6 on ward. These results suggested that MT-RRM approach can be more effective method for genomic prediction of WU.

**Figure 5:**
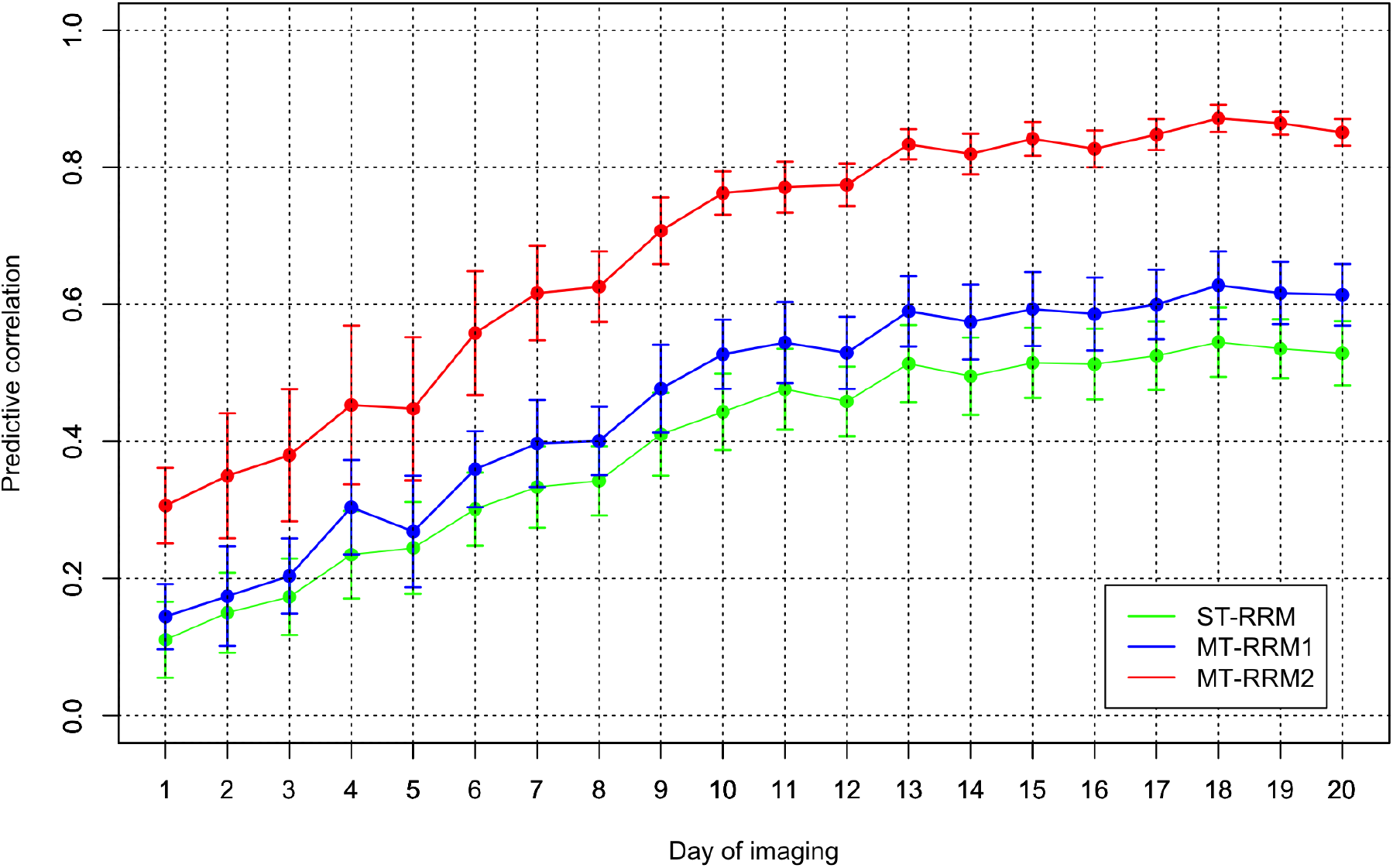
Pearson correlation of water use from cross-validation scenario 1. ST-RRM: single-trait random regression model; MT-RRM1: multi-trait random regression model using the water use and projected shoot area of training data; MT-RRM2: multi-trait random regression model using the water use and projected shoot area of training data as well as the PSA of testing data.

The objective of the second CV scenario was to evaluate the abilities of MT-RRM1 and MT-RRM2 to forecast genetic values at future time points using phenotypes recorded at earlier time points. The design of two MT-RRM were similar to those described above (Figure 1). The first MT-RRM, MT-RRM1, was fit using a training set with WU and PSA data collected from the first 10 time points, and was used to predict genetic values for WU for the subsequent time points. The second MT-RRM, MT-RRM2, was fit using PSA values for all 20 time points and WU values for the first 10 time points, and was used to predict genetic values for WU for the last 10 time points. Figure 6 shows the predictive correlation of WU for each of the models evaluated. The prediction accuracy for each method was relatively constant over all days. However, the prediction accuracy of MT-RRM were greatly higher than ST-RRM, indicating that inclusion of additional information from PSA can improve the ability to forecast WU. For ST-RRM, prediction accuracy ranged from 0.54 to 0.57, while values for MT-RRM1 and MT-RRM2 range from 0.79 to 0.83 and 0.84 to 0.91, respectively. Collectively, these results indicate that joint analysis of PSA and WU with the MT-RRM improves the ability to forecast future genetic values for WU.

**Figure 6:**
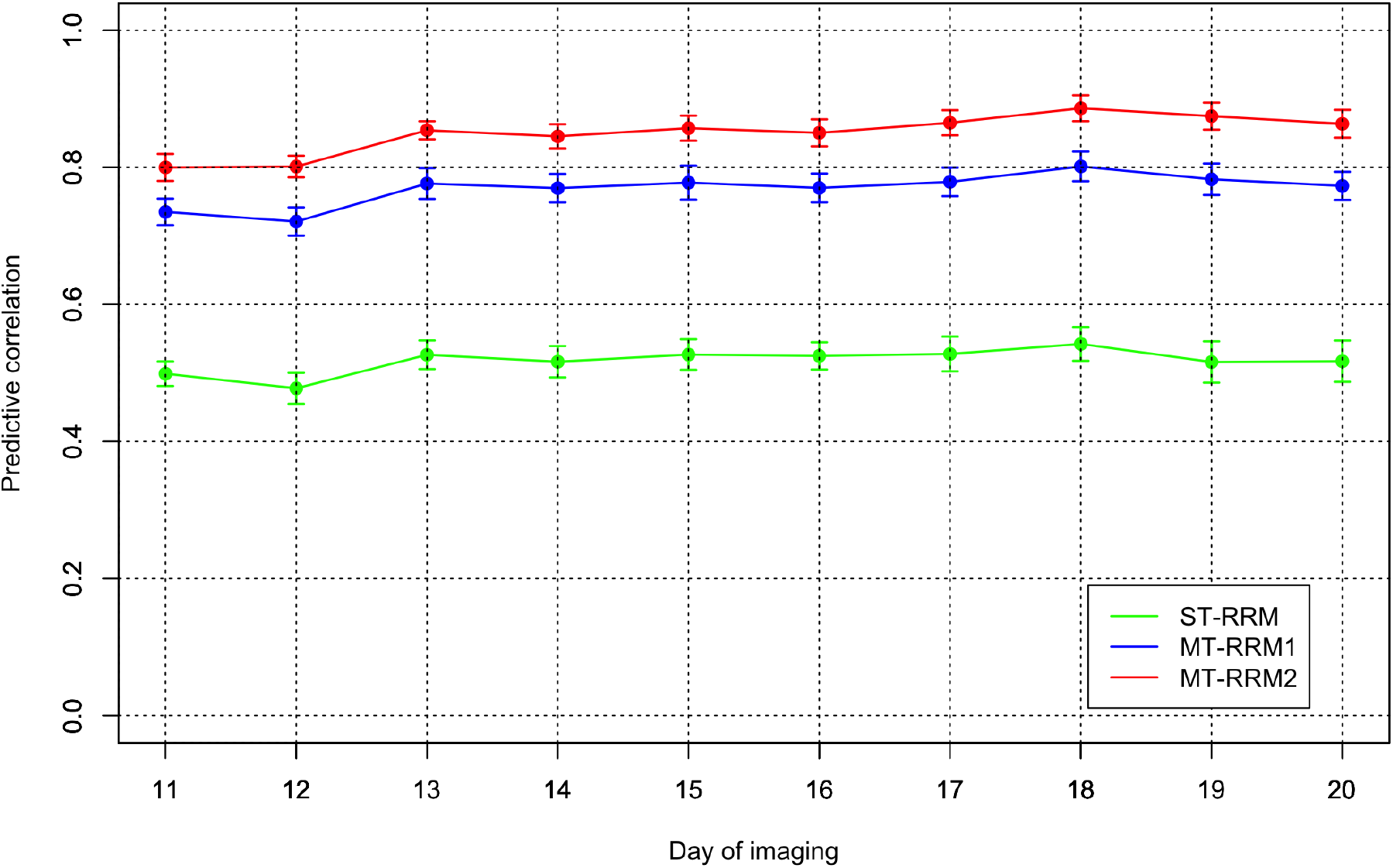
Pearson correlation of water use from cross-validation scenario 2. ST-RRM: single-trait random regression model; MT-RRM1: multi-trait random regression model using WU and PSA from 1 to 10 time periods in the training data; MT-RRM2: multi-trait random regression model using WU from 1 to 10 time periods and PSA at all the time periods in the training data.

## Discussion

Advances in HTP has provided plant breeders with a new suite of tools to assess morphological and physiological traits in a non-destructive manner for large populations at frequent time intervals throughout the growing season (Fahlgren et al., 2015). These platforms facilitate the collection of data that provide important insights into the morpho-physiological basis of complex traits. Thus, with these technologies complex traits such as drought tolerance can be decomposed into component traits to better understand the basis of these traits and improve the development of varieties with increased resilience (Berger et al., 2010). Although these platforms provide a powerful means to quantify complex traits in large populations, some physiological traits require specialized equipment or must be recorded during a specific time of day (e.g., transpiration or chlorophyll fluorescence) (Tardieu et al., 2017). Thus, in many cases these data may only be available for a subset of the population.

HTP is often used to record a number of traits on the same individuals. In some cases, physiological traits that are difficult to measure may be correlated with traits that are more accessible and can be recorded with greater ease. In such cases, MT genomic prediction frameworks provide an excellent solution to utilize partial records and predict genetic values for the physiological trait in individuals with missing data. Jia and Jannink (2012) demonstrated that MT models improve prediction accuracy particularly for traits with low heritability. In the current study, we utilized a MT approach in a RRM framework to predict genetic values for WU, a difficult to measure trait with low heritability, by joint analysis with PSA, which exhibits higher heritability and is easier to measure. Since WU shows a positive correlation with PSA, we hypothesized that the MT-RRM framework can improve predictions for WU.

## Genetic components of HTP image traits

Since WU is difficult to quantify directly in cereals such as rice, few studies have measured WU or water use efficiency, while most studies have sought to utilize indirect measurements of WU or water use efficiency for genetic analyses (This et al., 2010; Rebolledo et al., 2013; Feldman et al., 2018; Momen et al., 2019b). Consistent with the current study, Feldman et al. (2018), which utilized a HTP platform to quantify temporal water use and plant size in the C4 species *Setaria* grown in contrasting water regimes, reported moderate broad sense heritability (*H*^2^) values for WU, and higher *H*^2^ for plant size. Moreover, Feldman et al. (2018) showed that *H*^2^ varied throughout the experiment with lower *H*^2^ observed during the initial time points and higher *H*^2^ values observed during the middle time points. In our study, *h*^2^ values for WU in early time points were lower compared to those observed during the later time points. The plants in the current study were relatively small during the initial time points and therefore less amount of water is lost each day. Thus, water loss during these periods may be heavily influenced by environmental factors such as soil temperature or irradiation. Similar temporal trends have been reported for plant height in sorghum (Fernandes et al., 2018). Thus, given the moderate *h*^2^ values for WU and the temporal variability in *h*^2^, selection for this trait may be difficult in breeding programs. Conversely, *h*^2^ for PSA was relatively stable throughout the experiment, indicating that *h*^2^ for PSA may be less affected by temporal environmental effects compared to WU.

Multi-trait approaches are particularly advantageous when one target trait has low heritability and is correlated to a secondary non-target trait with higher heritability (Mrode, 2014). Joint analysis using a MT model can improve prediction of genetic values for low heritability trait and thus improve selection in plant breeding programs. In the current study, we showed a benefit of using MT-RRM for WU which had a positive genomic correlation with PSA. Thus, we proposed that joint analysis of WU with PSA can improve predictions of genetic values for WU. In a recent study, Momen et al. (2019b) examined the relationships between single time point measurements of WU, root biomass, water use efficiency, and PSA. According to the result, WU showed a moderate to strong positive correlation with PSA, root biomass, and water use efficiency, ranging from 0.48 to 0.85 (Momen et al., 2019b). Although we utilized PSA as the indicator trait in this study, it is expected that root biomass and water use efficiency can be leveraged for genomic prediction for WU using the MT.

## Predictive performance of MT-RRM

The MT-RRM framework offers several advantages over conventional single-trait genomic best linear unbiased prediction (ST-GBLUP) approaches. First, the random regression framework provides a tractable means to predict genetic values for temporal traits. The RRM uses covariances functions to model the genetic and environmental covariance between time points, and has been shown to improve prediction of genetic values compared to a ST-GBLUP approach (Campbell et al., 2018). Secondly, because the covariance function expresses the genetic covariance between time points using a continuous function, the RRM can be used to predict genetic values at time points with no records (Momen et al., 2019a). Thus, we can leverage the RRM framework to forecast future genetic values. Finally, as mentioned above, the joint analysis of MT can improve prediction accuracy for traits with low heritability. In the current study, we designed two CV to evaluate the ability to predict genetic values in unobserved accessions for a trait with lower heritability, and assessed the ability of the MT-RRM to predict future genetic values for accessions with records. Because the sample size and the number of time points used were relatively small, both ST- and MT-RRMs took less than 10 minutes to complete the longitudinal analyses on 64bit Linux with Intel Core i7-6950X (3.0GHz).

The first CV scenario was designed to evaluate the ability of MT-RRM to predict genetic values for WU in accessions without any records. Consistent with our expectations, MT-RRM had a better predictive ability than ST-RRM. The predictive ability of the MTRRM was further improved when PSA records were available for accessions in the testing population. The effectiveness of MT genomic models has been investigated extensively and have reported improved prediction accuracy compared to a ST model (Jia and Jannink, 2012; Guo et al., 2014; Okeke et al., 2017; Fernandes et al., 2018). For instance, Guo et al. (2014) compared prediction accuracy from ST- and MT-GBLUP using simulated data. MT-GBLUP showed better predictive performance when the target trait had lower heritability compared to the non-target trait and when the target trait had a greater number of missing observations (Guo et al., 2014). However, the majority of these studies have focused on traits recorded at a single time point. In the current study, we used a MT approach for prediction of bivariate traits with longitudinal records, and observed similar results. As suggested by the previous studies, an increase of prediction accuracy by MT-RRM in this study may result from a relative lower heritability of WU than PSA and the high degree of shared genetic signals with PSA (Momen et al., 2019b). The results of CV1 showed that prediction accuracies from all the models were more stable at later time periods, which is similar to the temporal trends in prediction accuracy observed for PSA reported by Momen et al. (2019a) that was obtained using a ST-RRM. The accuracy of genomic prediction largely depends on the heritability of the trait (Hayes et al., 2009). Thus, the lower predictions at the initial time points may be the result of the lower heritability observed during these periods. Moreover, early observations are recorded on seedlings that have just started to tiller. At this stage the plants may not have accumulated enough biomass and have low transpiration demands, to discern genotypic variation in water use from environmental variation.

Genomic predictions based on small number of records are a major concern in many practical applications, especially for a trait that is difficult or costly to measure because it can reduce phenotyping costs. As expected, the MT approaches (MT-RRM1 and MT-RRM2) in CV2 resulted in improvements compared to the ST-RRM with gains of 0.26 and 0.33, on average, for MT-RRM1 and MT-RRM2, respectively. Our results suggest that MT-RRM can be a powerful approach for forecasting future phenotypes using records from earlier periods. In this study, we examined prediction accuracies from 11 to 20 days in CV2. However, the trends in prediction accuracy were relatively stable across time points. Thus, forecasting based on records at further earlier time periods could be implemented without a loss of prediction accuracy as reported by Momen et al. (2019a). However, the performance of these forecasting approaches will likely be highly dependent on the genomic correlation between the time points used to train the prediction model and the time points in which predictions will be made. Lastly, it should be noted that the best prediction performance delivered by MT-RRM2 in both scenarios may be due to the fact that the training-testing sets partitioning is not completely independent in a strict sense. However, a situation akin to this occurs in practice and an approach such as MT-RRM2 would be still worthwhile to test.

We employed an unweighted two-stage approach to obtain adjusted means because of its simplicity and computational efficiency. However, a single-stage analysis is often considered as a more appropriate method to account for systematic effects due to heterogeneity of covariances among adjusted means (Möhring and Piepho, 2009; Piepho et al., 2012). Thus, we also explored a single-stage analysis by fitting all the systematic effects in RRM. We observed a high correlation (0.92) between the genetic values from the single-stage and the unweighted two-stage analyses across 20 time points. This is likely because the current dataset is obtained from the control condition in a greenhouse, which may yield a more homogeneous variance-covariance structure of errors between adjusted means compared to heterogeneous data typically collected from multi environment field trials. A weighted two-stage approach (Smith et al., 2001; Piepho et al., 2012) was not considered in the current study because of the limitation of the GIBBS3F90 program to perform such an analysis.

## Conclusion

To our knowledge, this is the first study that applied the MT-RRM to HTP-derived temporal traits in plants. We demonstrated that MT-RRM is a robust and flexible approach that can be used to improve prediction accuracy for a trait with a limited number of records or low heritability. Thus, in the case of breeding for morpho-physiological traits, the MT-RRM can improve prediction accuracy for physiological traits that may have low heritability or are difficult to measure in large populations.

## Acknowledgments

This work was supported by the National Science Foundation under Grant Number 1736192 to HW and GM, and Virginia Polytechnic Institute and State University startup funds to GM.

## Author contribution statement

This work was supported by the National Science Foundation under Grant Number 1736192 to HW and GM, and Virginia Polytechnic Institute and State University startup funds to GM. MTC and HW designed and conducted the experiments. TB and MM analyzed the data. TB and GM conceived the idea and wrote the manuscript. MTC, MM and HW discussed results and revised the manuscript. GM supervised and directed the study. All authors read and approved the manuscript.

